# The evolution of the 9aaTAD domain in Sp2 proteins: inactivation with valines and intron reservoirs

**DOI:** 10.1101/570598

**Authors:** Martin Piskacek, Marek Havelka, Kristina Jendruchova, Andrea Knight, Liam P. Keegan

## Abstract

The Sp1 transcription factor has been defined as glutamine-rich activator. The Nine amino acid TransActivation Domains (9aaTAD) have been identified in numerous transcription activators. Here, we identified the conserved 9aaTAD motif in the Sp1 and in all nine members of SP family with broad natural 9aaTAD variations. We showed by the amino acid substitutions that the glutamine residues are completely dispensable for 9aaTADs function. We described the 9aaTAD domains’ origin and evolutionary history. The ancestral Sp2 gene with inactive 9aaTAD has duplicated in early chordates and created new paralogs Sp1, Sp3 and Sp4. We discovered that the accumulation of valines in the 9aaTADs correlated with the domain inactivation. The Sp2 activation domain, whose dormancy have lasted over 100 million years during chordate evolution, enabled later diversification in the Sp1-4 clade, including both repressors and activators. The new paralogs Sp1 and Sp3 activation domains have regained their original activator function by loss of valines in their 9aaTADs.

## INTRODUCTION

Our longstanding effort has been to determine the pattern of trans-activating sequences in the eukaryotic activators. Previously, we reported successful identifications of the Nine amino acid TransActivation Domains, 9aaTADs, in a large set of the transcription activators that universally recruit multiple mediators of transcription (1–11).

We and others have shown that the activation domains have the competence to activate transcription as small peptides (above 9 to 14 amino acids long) in over 40 activators including Gal4, E2A, MLL and p53 (12–17). Although the activation domains have enormous variability, they are universally recognized by transcriptional machinery throughout eukaryotes (18). Currently, all our tested human 9aaTAD activation domains were shown to be functional in yeast. Therefore we considered universal function of 9aaTAD activation domains in eukaryotes as a further associated property of the 9aaTAD family (12, 14, 18, 19). In addition to the amino acid pattern and universal function in eukaryotes, the 9aaTAD activation domains are further characterized by tandem hydrophobic clusters separated by a hydrophilic region (12, 14). The 9aaTAD domains are well balanced with hydrophilic amino acids, which might be either positively or negatively charged. From the structural data for the E2A and MLL activation domains in complex with the KIX domain, we observed that 9aaTAD domains formed short helices whose length varies from 9 to 12 aa (13). Online 9aaTAD prediction (using a residue position matrix search) is available on www.piskacek.org. The curated 9aaTAD activation domains had been annotated on protein database UniProt (by database search for 9aaTAD).

Earlier, the activation domains have been categorised and their functions were linked to over-represented amino acids (20). The aspartate activation domains were described as acidic (21) or glutamine-rich (22, 23) or proline-rich activation domains (24, 25). Accordingly, the Gcn4, Gal4 and p53 activation domains have been designated as acidic activators (26–29). The next studies revealed some controversies and demonstrated the importance of the hydrophobic amino acids for Gcn4 (30–33) and p53 activators (8, 34–40). Similarly, in the original report for the glutamine-rich activator Sp1(41), the importance of hydrophobic amino acids was already recognized, regardless of the overrepresented glutamines. Nevertheless, the designation as acidic or glutamine-rich activation domains persists to date (42–49). In the human genome, there are about 1,691 annotated transcription factors, divided into 68 groups according to their conserved DNA binding domain (50). The C2H2-type zinc finger is the largest group (682 members), which includes the evolutionally conserved family of KLF activators (Krüppel-like factors). The SP activators (Specificity proteins Sp1-9) belong to a new evolutional branch derived from the ancestral KLF family. The SP activators have been linked to cell proliferation, embryonic development, tissue differentiation, Wnt signaling pathway, metabolism, and their dysregulation has been implicated in a number of human diseases and cancers (51–56). This report has focused on the SP family and followed the 9aaTAD domain’s evolution.

## MATERIALS AND METHODS

### Constructs

The construct pBTM116-HA was generated by an insertion of the HA cassette into the *Eco*RI site of the vector pBTM116. The constructs G1-G45 and H1-H45 were generated by PCR and sub-cloned into pBTM116 *Eco*RI and *Bam*HI sites. All constructs have a spacer of three amino acids inserted into the *Eco*RI site; peptide GSG. All constructs have been sequenced by Eurofins Genomics. Further detailed information about constructs and primer sequences are available on request.

### Assessment of enzyme activities

The β-galactosidase activity was determined in the yeast strain L40 (57, 58). The strain L40 has a chromosomally integrated *lacZ* reporter driven by the *lexA* operator. In all hybrid assays, we used 2μ vector pBTM116 for generation of the LexA hybrids. The yeast strain L40, *Saccharomyces cerevisiae* Genotype: *MATa ade2 his3 leu2 trp1 LYS::lexA-HIS3 URA3::lexA-LacZ*, is deposited at ATCC (#MYA-3332). The average value of the β-galactosidase activities from two independent transformants is presented as a percentage of the reference with the standard deviation (means and plusmn; SD; n = 2). We standardized all results to the previously reported Gal4 construct HaY including the 9aaTAD domain with the activity set to 100% (14).

### Databases used in the study

UniProt, ExPASy, NCBI, KEGG http://www.genome.jp, Japanese Lamprey Genome Project http://jlampreygenome.imcb.a-star.edu.sg/blast/, Sun Yat-Sen University Lancelet Genome Project http://genome.bucm.edu.cn/lancelet/blast.php, Compagen Genomics Platform http://www.compagen.org/blast.html, Florida University Neurobase https://neurobase.rc.ufl.edu/, UCSC genomic annotation https://genome.ucsc.edu.

## RESULTS

### The 9aaTAD activation domains occur in proteins previously used to define acidic, glutamine-rich (Sp1) or proline-rich activator classes

The Gal4 activation domains have been reported as acidic (aspartic acid-rich), the Sp1 activation domain as glutamine-rich, and the NFIC activation domains as proline-rich (21, 25, 41, 44). However, the importance of the acidic aspartic residues for Gal4 function addressed already by the Johnston group (59). Similarly for Sp1, the authors have demonstrated a greater importance of hydrophobic amino acids, rather than the overrepresented glutamines (41).

Previously, we have identified the 9aaTAD activation domains in the Gal4 (17) and in p53 acidic activators (14). In this study, we aimed to determine the 9aaTAD presence in other members of the SP family, where the 9aaTAD position would correspond with the Sp1 region 467-472 that has been shown to be essential for transcriptional activation (41). Using our online 9aaTAD prediction, we obtained two positive prediction hits for Sp2 (using the most stringent pattern and with 83% and 92% match for the 9aaTAD prediction). Most importantly, the latter match (sequence GEVQTVLVQ in the Sp2 region 361-369) corresponds with the glutamine-rich activation domain of Sp1 (region 467-472).

We have then generated LexA constructs, which included the DNA binding domain LexA and only the 9aaTAD from all nine SP activators and tested them for the ability to activate transcription (Figure 1 and 2). The predicted Sp1 activation domain includes glutamines, which are not conserved in Sp5, 6, 7, 8, 9 (Figure 2) and are not generally conserved in other 9aaTAD domains. Similarly, the acidic activation domain of the Gal4 activator includes acidic aspartic acids, which are not generally conserved in other 9aaTAD domains (12, 14, 17). In order to prove that the function of the 9aaTADs was not solely dependent on overrepresented aspartates nor on glutamines (which are present in the Gal4 and Sp1 activation domains), we have substituted all Gal4 aspartic residues for glutamines (construct Gal4-Q, non-acidic Gal4 9aaTAD) and all Sp1 glutamines and asparagine for aspartic acid residues (construct Sp1-Ac, acidic Sp1 9aaTAD), and showed their ability to activate transcription (Figure 1).

By using our online 9aaTAD prediction, we have searched for the 9aaTADs in the proline-rich activation domain of the NFIC / CTF activator (region 388-487). We have identified two overlapping 9aaTAD motifs (sequences DPLK DLVSL and KDLV SLACD with 83% and 92% match for the 9aaTAD prediction) in the NFIC proline-rich region (the pattern for clusters of less stringent repeats was used). Moreover, we also identified both corresponding 9aaTAD motifs in the very distal orthologs <u>Sko</u>NFI (from <u>*S*</u>*accoglossus* <u>*ko*</u>*walevskii*, acorn worm), and <u>cin</u>NFI (from <u>*C*</u>*iona* <u>*in*</u>*testinalis*, sea squirt). Furthermore, in the NF family, prolines are not conserved in any position of the human paralogs (NFA, NFB, NFIC, NFIX), nor in the <u>Sko</u>NFI or cinNFI orthologs, however the 9aaTADs were well conserved in all of them (Figure 1). We have generated a LexA construct, which included only the 9aaTAD from the NFIC activator, and showed its ability to activate transcription (Figure 1).

**Figure 1.**
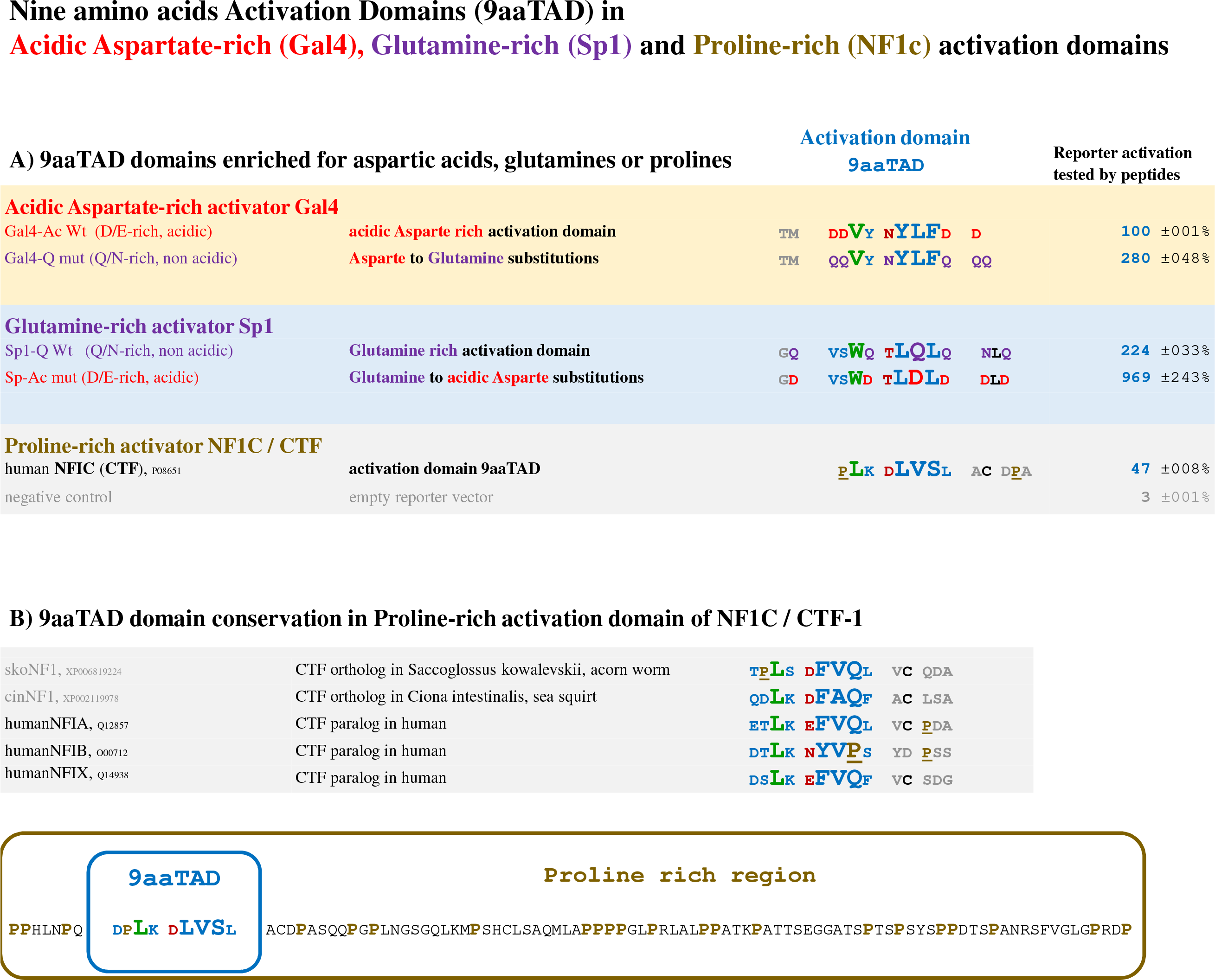
The activation domains subfamilies. A) The regions with identified activation domains 9aaTAD in Gal4 (in red), Sp1 (in purpura), their mutants, and NF1C / CTF (in sand brown) were tested in reporter assay with hybrid LexA DNA binding domain for the capacity to activate transcription. The mutants of Gal4 and Sp1 were designed to substitute aspartates (in red) for glutamines (in purpura) and vice versa. B) The NF1 / CTF activator, orthologs and paralogs are shown to demonstrate 9aaTAD motif conservation in their kin (adjacent conserved cysteines are in black). At the bottom of the figure, the entire region of previously reported proline-rich activation domain of the NF1 / CTF is shown; the activation domain 9aaTAD is marked (prolines in sand brown). The activation domains 9aaTAD are coloured for fast orientation. The LexA-Gal4 hybrid constructs assayed in L40 strain for transactivation activity are shown.

### The 9aaTAD activation domain in the human SP family

As described above, we have identified the 9aaTAD motif in the human transcription factor Sp2 and consequently, we have aligned all human SP paralogs and found conservation of the 9aaTAD motif in all nine members of the family (Figure 2).

The human SP transcription factors share sequence homology in their activation domains and could be divided into two clades (Figure 2); i) the Sp1-4 clade with a single tryptophan (single letter code W) in the activation domain (41), excluding the Sp2, whose activation domain lacks tryptophan, and ii) the Sp6-9 clade with two tryptophans in the activation domain, excluding Sp5, the sequence of which differs significantly from other members of Sp6-9 clade.

**Figure 2.**
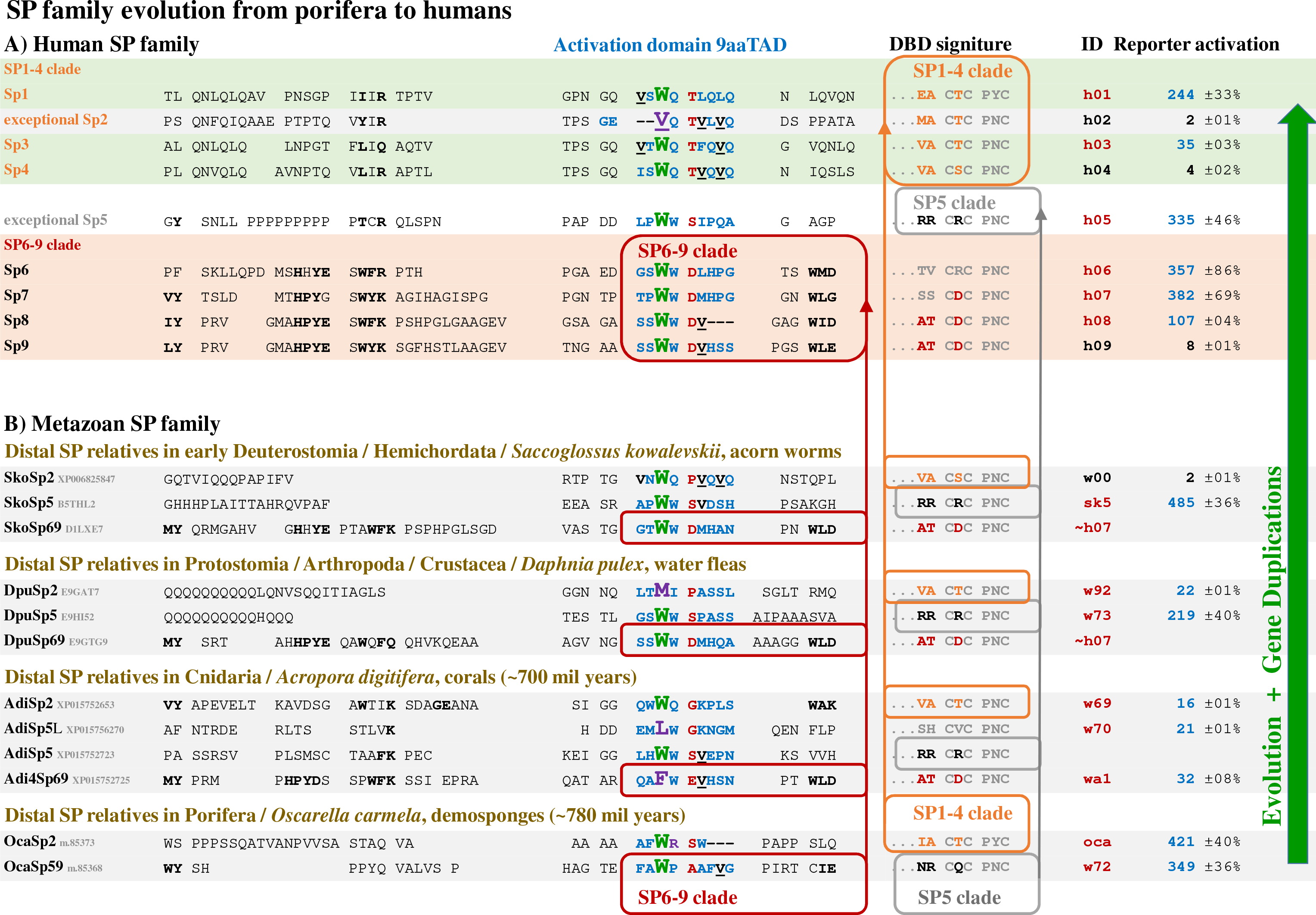
The SP family. A) The 9aaTAD activation domains identified in the SP family are shown for humans, for early deuterostomes / hemichordates / *Saccoglossus kowalevskii* (acorn worms), for protostomes / arthropods / crustacea / *Daphnia pulex* (water fleas), for cnidarians / *Acropora digitifera* (corals), and for poriferans / *Oscarella carmela* (demosponges). The designation *Sp69* reflected uncertainty of origin (a single member of the SP6-9 clade in those animals), whereas designation *Sp59* in *Oscarella carmela* pointed to common ancestor of the latter *Sp5* and Sp6-9 clade. The peptides with capacity to activate transcription are with red ID and those without are in black (five-fold induction below the threshold). The activation domains with similarity are referred to by ~ marks. A link of the *Sp5 genes* and *Sp6-9 genes* is marked in grey boxes and *Sp2 genes* in yellow boxes. The most intriguing variations are in purpura. The dash symbol means amino acid deletion in human *Sp8*. The activation domains 9aaTAD are coloured for fast orientation. Valines in the 9aaTAD are in black and underlined (the pattern V-Q/H correlated here with the deactivation of the 9aaTAD activation domain in <u>*h*</u>*Sp2*, <u>*h*</u>*Sp4*, <u>*h*</u>*Sp9* and <u>*Sko*</u>*Sp2*).

We have generated LexA constructs h01 - h09, which included only the 9aaTADs from Sp1 to Sp9 proteins, and tested them for the ability to activate transcription (Figure 2). The members of the SP family were reported to be activators of transcription (52), with the exception of Sp2, which is a weak activator with little or no capacity for additive or synergistic transactivation (60–66), and functions as a repressor (67). Apart from the h02, h04 and h09, all other constructs h01, h03, h05, h06, h07 and h08 showed strong capacity to activate transcription (Figure 2). The deactivated 9aaTADs correlated with the accumulation of valines in the Sp2, Sp4 and Sp9 activation domains and follow a V-Q/H amino acid pattern. Similarly, we found an inactive Sp7 activation domain in the spider mite (*Tetranychus urticae*), where the 9aaTAD deactivation correlated with valine acquisition and followed the V-Q/H amino acid pattern (**Suppl. Figure S1**). We could also identify a 9aaTAD deletion instead of deactivation (**Suppl. Figure S2**). The prime aim of this study was the identification of functional activation domains. Those which were unable to activate transcription under our experimental conditions are also reported, but without further investigation. Nevertheless, the expression of inactive constructs was monitored by Western blotting (**Suppl. Figure S3**). Of note, in this study, we did not test the entire proteins but only the distinct regions predicted to have activation function. For example, the 9aaTAD from human Sp2 failed to activate transcription in our assay. Our result is compliant with previous reports, finding Sp2 a weak activator with little or no capacity for additive or synergistic transactivation (65). The Sp2 was reported to specifically bind DNA, occupies distal enhancer elements (68–70) and functions as repressor (67). Differently, human Sp1 and Sp3 were reported as activators of transcription (71–73).

### Evolution of the 9aaTAD activation domain in SP proteins began in Sea Sponges about 780mil years ago

The SP family is a new branch derived from the KLF family. The ancestral *SP genes* appear in demosponges ~800 million years ago (Mya) (74, 75). The SP family conservation is remarkably strong, particularly in the region of the DNA binding domains. We have followed the diversification of activation domains in SP family proteins in both vertebrates and invertebrates (**Suppl. Figures S4 and S5**) and showed diversification in early-branching metazoan clades Sp1-4 (**Suppl. Figure S6A**), Sp6-9 (**Suppl. Figure S6B**), Sp5 (**Suppl. Figure S6C**) and Sp2 (Figure 3) in the following paragraphs. Using blast searches (see list of Databases used in Materials and Methods), we have identified the 9aaTAD activation domains in human SP activators and their distal ancestors, including in the hemichordate *Saccoglossus kowalevskii* (acorn worm, early deuterostomes), in the crustacean *Daphnia pulex* (water flea, arthropods; branch of ecdysozoans, protostomes), in the cnidarian *Acropora digitifera* (coral; branch of eumetazoans about 700 Mya), in the poriferan *Oscarella carmela* and *Amphimedon queenslandica* (demosponge; early branch of metazoan about 780 Mya) (74) (Figure 2).

**Figure 3.**
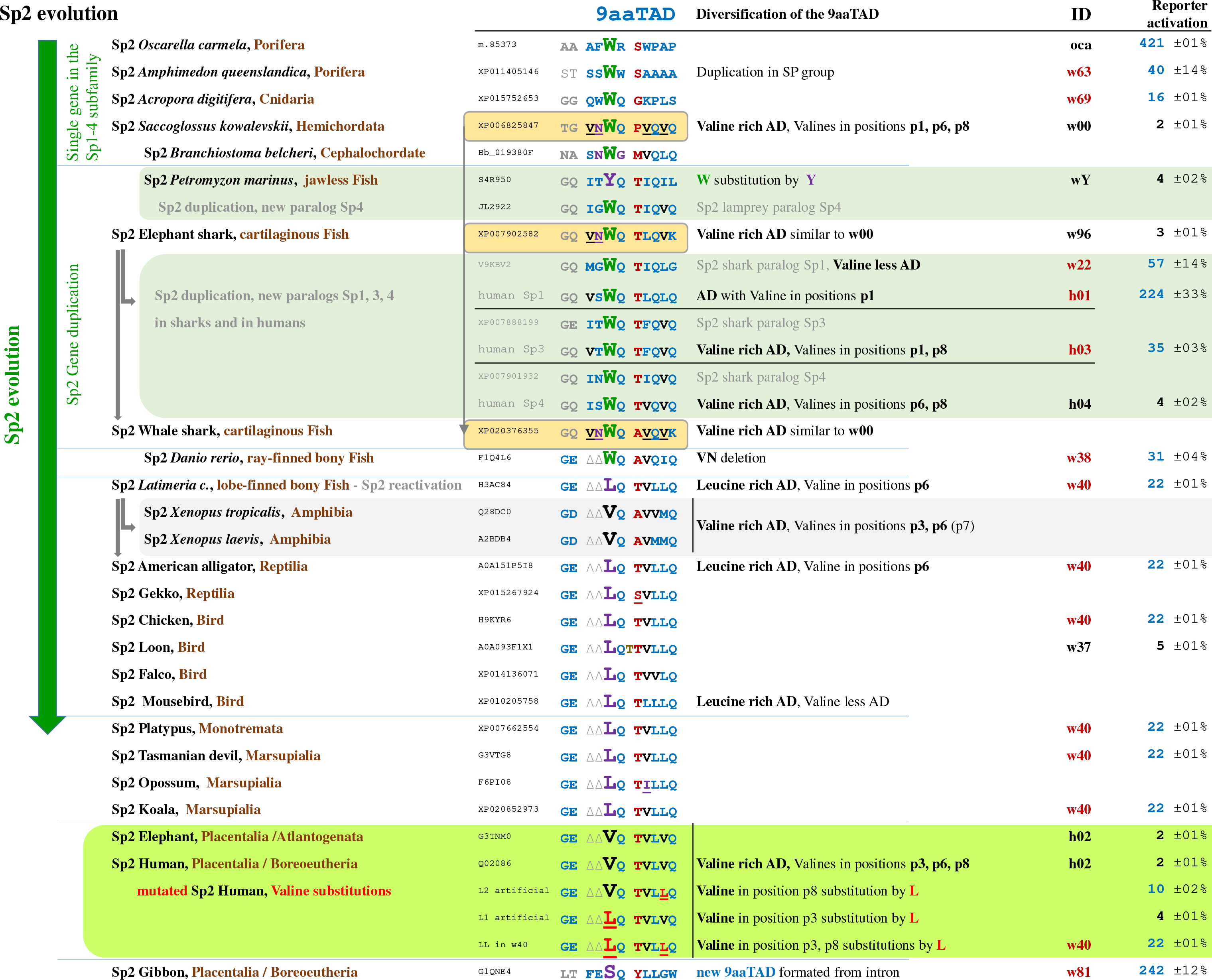
SP2 clade evolution. The 9aaTAD activation domains identified by similarity (BLAST search in metazoans) in the SP2 clade from early branched metazoans to humans are shown. Some of them were tested in a reporter assay with a hybrid LexA DNA binding domain for the capacity to activate transcription. The peptide IDs with capacity to activate transcription are in red and those without are in black (5-fold induction above threshold). The most intriguing variations are in purpura and black. The paralogs (originated from *Sp2 gene* duplication in elephant shark) are in grey and the *Sp2* paralogs of cartilaginous and hemichordates, which show high sequence similarity are in grey boxes. The 9aaTAD activation domains are coloured for fast orientation. Valines in the 9aaTADs are in black. Valine rich (V1 in position p1, V2 in position p6, V3 in position p8) activation domains (AD) with tryptophan (in position p3) are in hemichordates and cartilaginous fish. A link of the first *Sp1-4 genes* with the *Sp2* orthologs and paralogs (in grey) is shown (similarity of hemichordates and cartilaginous fish Sp2 9aaTAD domains are underlined). Diversified branches are indicated.

We generated LexA constructs for the activation domains of ancestral *SP genes*, designated as <u>*Sko*</u>Sp2 and <u>*Sko*</u>Sp5 (<u>*S*</u>*accoglossus* <u>*ko*</u>*walevskii*); <u>*Dpu*</u>*Sp2* and <u>*Dpu*</u>*Sp5* (<u>*D*</u>*aphnia* <u>*pu*</u>*lex*); <u>*Adi*</u>*Sp2*, <u>*Adi*</u>*Sp5* and <u>*Adi*</u>*Sp69* (<u>*A*</u>*cropora* <u>*di*</u>*gitifera*); and <u>*Oca*</u>*Sp2*, *OscaSp59* (<u>*O*</u>*scarella* <u>*ca*</u>*rmela*) and tested them for the ability to activate transcription (Figure 2) (the designation *Sp69* and *Sp59* reflects uncertainty of origin). <u>*Sko*</u>*Sp5* and <u>*Dpu*</u>*Sp5* genes are almost identical to human *Sp7*, and therefore omitted. We tested and confirmed that most of constructs showed the capacity to activate transcription. The 9aaTAD of the *Saccoglossus* <u>*Sko*</u>*Sp2* (construct w00, with accumulated valines within the 9aaTAD domain) was fully deactivated (Figure 2).

### Conservation of the 9aaTADs in the SP family proteins in vertebrates

Next, we focused on the evolution of the SP family in vertebrates. As representative examples for SP family evolution in vertebrates, we selected the cartilaginous fish *Callorhinchus milii* (elephant shark), ray-finned fish *Danio rerio* (zebrafish), lobe-finned fish *Latimeria chalumnae* (coelacanth), and bird *Gallus gallus* (progenitor of the domestic chicken), and compared their 9aaTAD activation domains with the human ones (**Suppl. Figure S5**). Most surprisingly, we have found a remarkably strong family conservation of the 9aaTADs in *Sp1, Sp3, Sp4, Sp5, Sp6, Sp7, Sp8* and *Sp9*, but not in *Sp2*. Furthermore, the *Sp7 gene* was missing altogether in *Gallus gallus*, but not in all birds. The missing *Gallus Sp7* was substituted by the *Pseudopodoces humilis* ortholog in **Suppl. Figure S5.** The summary of the bird *Sp7 gene*s, which are missing in some but not all birds, are shown in **Suppl. Table 1**.

Throughout evolution, the loss of the *Sp7 gene* has occurred in several bird clades. Importantly, *Sp7* was lost only in some passeriformes. The *Sp7*-less passeriformes members (crow, flycatcher and finch) could have not been the ancestral for other *Sp7*-less clades e.g. falconiformes (76). Therefore, the loss of *Sp7* gene was not a single event during the bird evolution, suggesting a genetic selection in the phylogenetic distribution, rather than simple random events.

### The SP1-4 clade diversification

The SP1-4 clade found in early-branching metazoans (**Suppl. Figure S6A**) showed the same preference for a single tryptophan in position p3 of the 9aaTAD activation domains as in vertebrates (**Suppl. Figures S5**). However, there are also exceptions, with twin tryptophan in arthropods, *Strigamia maritima* (Leach) which is typical for the SP6-9 clade (Figure 2) and a tryptophan-free activation domain in molluscs *Biomphalaria glabrata*. Thus, the natural occurring 9aaTAD variance has shown that tryptophans, which dominated in the SP family, are not essential for 9aaTAD function. Similar observation was done for the 9aaTAD activation domain in p53 (14).

### The SP6-9 clade diversification

The *Sp69 gene* (the designation *Sp69* reflects uncertainty of origin, Figures 2) has already emerged in radiates and their 9aaTADs have two tryptophans in positions p3 and p4, which are well conserved in vertebrates (**Suppl. Figure S5**). The hemichordate *Saccoglossus kowalevskii* (acon worm) has been the only solitary member of the SP6-9 clade, with the <u>Sko</u>S*p69 gene*, which gave rise to the SP6-9 clade (gene duplication). Thus, *Callorhinchus milii* (elephant shark) already has four Sp69 paralogs, including the <u>*Cmi*</u>Sp6-9 genes. Similarly, the <u>Sko</u>*Sp2 gene* is a solitary member of the SP1-4 clade in the *Saccoglossus kowalevskii*. Higher in evolution, *Callorhinchus milii* already has four Sp2 paralogs including <u>Cmi</u>Sp1-4 (Phylogenetic tree, **Suppl. Figure S4**).

### The Sp5 evolution

From the sequence similarity and from the phylogenetic tree (Figure 2 **Suppl. Figure S4**), we predicted that the SP5 clade has emerged already in the earliest metazoans, before the SP6-9 clade (Figure 2). The DNA binding domain signature in the poriferan <u>*O*</u>scarella <u>ca</u>rmela <u>Oca</u>Sp59 (<u>**NR**</u>C<u>**Q**</u>-CPNC, designation *Sp59* in *Oscarella carmela* pointed to a common ancestor of the latter *Sp5* and *Sp6-9* clades) best matches the cnidarian <u>*A*</u>cropora <u>di</u>gitifera <u>Adi</u>Sp5 (<u>**RR**</u>C<u>**R**</u>-CPNC), the protostome <u>*D*</u>*aphnia* <u>*pu*</u>*lex* <u>*Dpu*</u>*Sp5* (<u>**RR**</u>C<u>**R**</u>-CPNC), the hemichordate <u>*S*</u>*accoglossus* <u>ko</u>walevskii <u>Sko</u>Sp5 (<u>**RR**</u>C<u>**R**</u>-CPNC) and is conserved up to <u>h</u>uman <u>*h*</u>*Sp5* (<u>**RR**</u>C<u>**R**</u>-CPNC). However, it less matches the diverged members of the SP6-9 clade; <u>*Adi*</u>*Sp9* (<u>AT</u>C<u>D</u>-CPNC) or <u>*h*</u>*Sp6* (<u>TV</u>C<u>**R**</u>-CPNC), <u>*h*</u>*Sp7* (<u>SS</u>C<u>D</u>**-**CPNC), <u>*h*</u>*Sp8* (<u>AT</u>C<u>D</u>**-**CPNC), and <u>*h*</u>*Sp9* (<u>AT</u>C<u>D</u>**-**CPNC) (Figure 2). The members of the SP5 clade also showed a preference for two tryptophans in positions p3 and p4 of their activation domains. The *Sp5* and *Sp69 genes* already largely diverged in already in cnidarian (*Acropora digitifera*; corals) and formed a separate clades. In protostomes, the SP5 clade shows the largest diversity we have ever observed for the 9aaTAD activation domains (**Suppl. Figure S6C**). We have generated constructs for selected members of the SP1-4 **(Suppl. Figure S6A)**, SP6-9 **(Suppl. Figures S6B)** and SP5 clades **(Suppl. Figures S6C)**. All tested constructs have the capacity to activate transcription.

### The Sp2 evolution

Next, we have generated constructs shown in (Figure 3) for the Sp2 activation domains including the w63 for *Amphimedon queenslandica* (demosponge, poriferan), constructs w69 for *Acropora digitifera* (coral, cnidarian), w00 for *Saccoglossus kowalevskii* (acorn worm, hemichordate), construct w96 for *Callorhinchus milii* (elephant shark, cartilaginous fish), construct w38 for *Danio rerio* (zebrafish, ray-finned bony fish), w40 for activation domains that are common for *Latimeria chalumnae* (coelacanth, lobe-finned bony fish), to American alligator and to gecko (reptiles), to *Gallus gallus* and egret (birds), to platypus (monotremates) and to Tasmanian devil, opossum and koala (marsupials), construct w37 for exceptional loon Sp2 activation domain (birds), construct h02 for Sp2 activation domains common to elephants and humans (representing two separate clades of placental mammals: atlantogenates and boreoeutheries) and construct w81 for the exceptional gibbon Sp2 activation domain (primates). Furthermore, we generated constructs w22 for the Sp1 activation domain 9aaTAD of *Callorhinchus milii*, which is an early diversified *Sp2* paralog in the SP1-4 clade.

Interestingly, not all tested Sp2 constructs showed the capacity to activate transcription. There are periods in metazoan evolution (from hemichordates to chondrichthyes and from early placental mammals to humans), in which the Sp2 activation domains have been deactivated. The deactivation of the 9aaTADs correlated with the accumulation of valines in their activation domains and followed an amino acid pattern of V-Q/N/K. Although tryptophans dominated in the SP activation domains (Figures 2), we have also found a tryptophan-less Sp2 activation domain in the sarcopterygians (in the lobe-finned bony fish *Latimeria chalumnae*) and in all their successors (higher vertebrates) (**Suppl. Figure S5**). We have then generated alignments and a simplified phylogenetic tree for *Saccoglossus* <u>*Sko*</u>*Sp2*, <u>*R*</u>*hincodon* <u>*ty*</u>*pus* <u>*Rty*</u>*Sp2* and <u>*C*</u>*allorhinchus* <u>*mi*</u>*lii* <u>*Cmi*</u>*Sp2* (Figures 2). The close orthologs of <u>*Sko*</u>*Sp2* seem to be <u>*Cmi*</u>*Sp2* and <u>*Rty*</u>*Sp2*, which also correlated with close similarities in their activation domains (**Suppl. Figure S4**).

In summary, we conclude that *Saccoglossus* <u>*Sko*</u>*Sp2* is the ancestral gene for the clade SP1-4. In chondrichthyes (cartilaginous fish) after *Sp2 gene* duplications, Sp1 regained activator function by *de-valinisation* (substitution of valines in the 9aaTAD for other amino acids), which remained during evolution all the way to humans. The *Sp2* has regained some activator function later in evolution (in sarcopterygians), but only up to the marsupials, not to the placentals (Figure 3).

From the phylogenetic tree (**Suppl. Figures S4)**, we have anticipated that the *Sp2 gene* originated directly from one of two ancestral *SP genes* found already in demosponges (Sp2 and Sp59; demosponges belong to early branched poriferan branched around ~800 Mya) **(Figure 2)**. Also, in the hemichordate *Saccoglossus kowalevskii* (acorn worm, belonging to early branched deuterostome, ~670 Mya), the *Sp2 gene* is the solitary member of the SP1-4 clade. From above, the *Sp2 gene* duplication gave rise to the SP1-4 clade found in jawed cartilaginous fish (chondrichthyes) such as *Callorhinchus milii* (elephant shark) and *Rhincodon typus* (Whale shark). The activation domain of this <u>*Sko*</u>*Sp2* gene with the sequence <u>VNWQ</u>-PV<u>QV</u>Q sustained conserved nearly fully in the *Rhincodon* <u>*Rty*</u>*Sp2* activation domain (<u>VNWQ</u>-A<u>VQV</u>K i.e. 7 identical and 8 similar residues) and almost in *Callorhinchus milii* <u>*Cmi*</u>*Sp2* activation domain with the sequence <u>VNWQ</u>-TL<u>QV</u>K. The activation domain of this <u>*Sko*</u>Sp2 gene (as well as the entire gene, phylogenetic tree **Suppl. Figures S4**) is less similar to other genes of the *Callorhinchus milii Sp1-4* clade. The <u>*Sko*</u>*Sp2*, <u>Cmi</u>Sp2 and <u>Rty</u>Sp2 have accumulated valines in their activation domains, which are then associated with the loss of their transcription activating function. After gene duplication, these valines were substituted by other amino acids (*de-valinisation*) in the activation domains of <u>*Cmi*</u>Sp1, which therefore regained their activation function. However, the <u>*Cmi*</u>*Sp2* remained deactivated (Figure 5).

**Figure 4.**
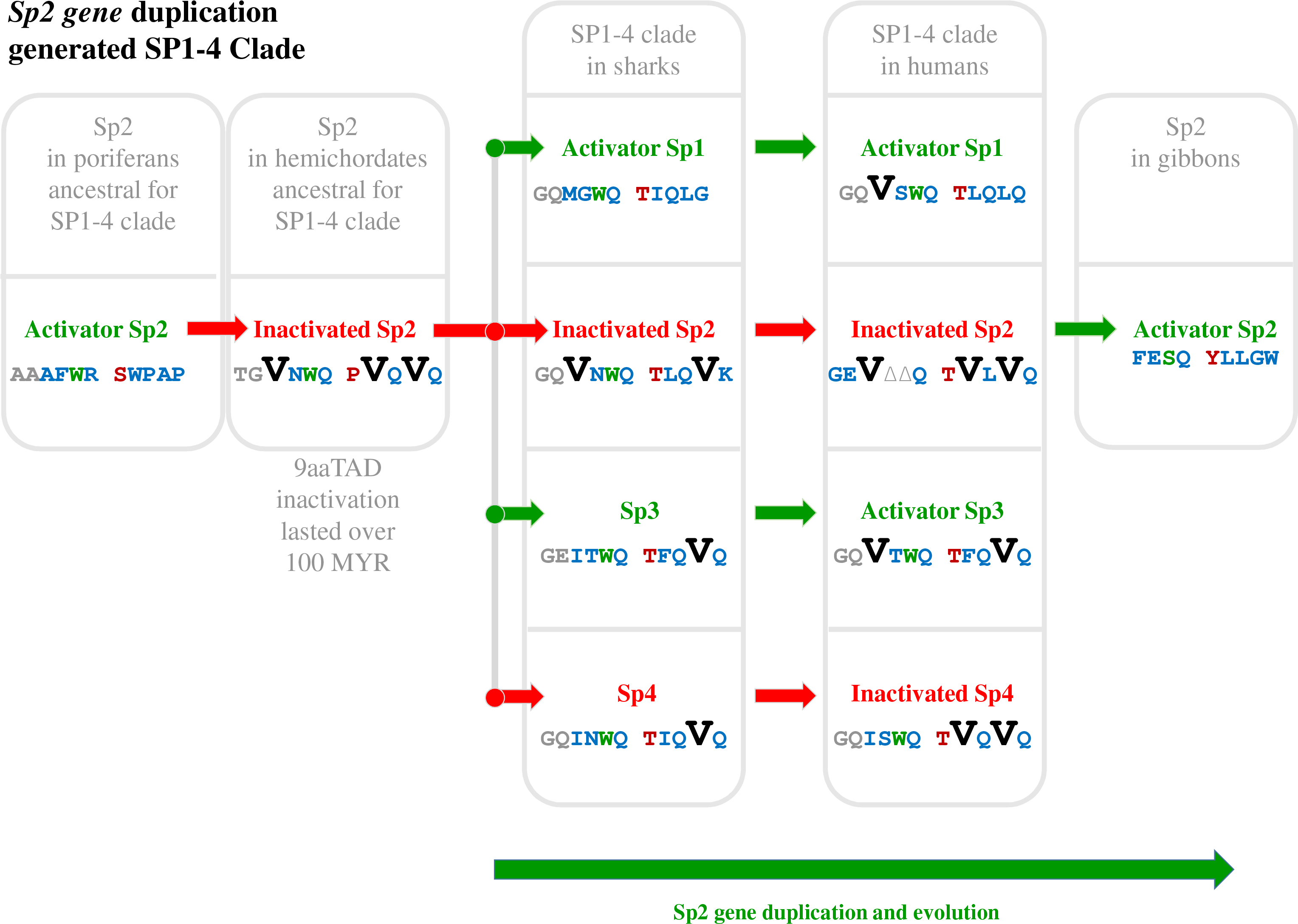
Schema of the SP1-4 clade evolution.

**Figure 5.**
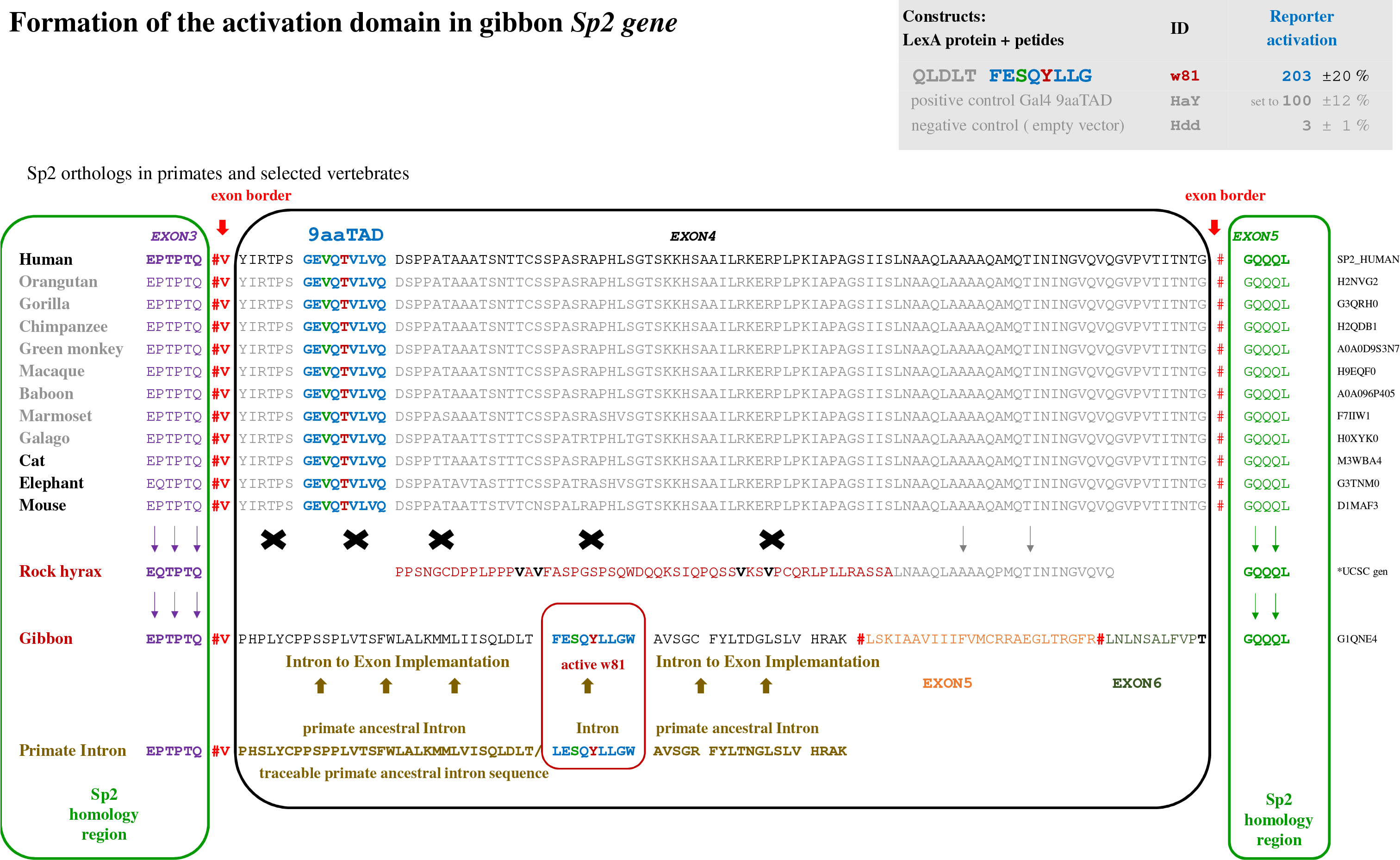
Reformation of the activation domain in the gibbon *Sp2 gene*. The Sp2 activation domain coded by *exon 4* conserved in placental mammals (primates, cats, elephants and mice are shown in grey) was lost in hyrax and gibbon. Their new exons originated from their ancestral introns (in sand brown, exon junction in red, *exon 3* in purpura, *exon 5* in green). This repair took placed on the border of the *exon 3* and *exon 4*, exactly as in takifugu and tetraodon *Sp2b genes* and crane *Sp2 gene*. In gibbon’s case, the intron provided a new 9aaTAD activation domain and their *Sp2 genes* render activator function. The missing gibbon ancestral intron was substituted in our figure by the human intron, which is close to all primates and should substitute for the gibbon ancestral intron well. Slashes denote frameshifts in intron sequence. The activation domains 9aaTAD are coloured for fast orientation. The exon junctions are marked by a red hashtag. *UCSC genomic annotation for Procavia capensis (rock hyrax) *Sp2 gene* in UCSC scaffold_25:102095-130179. We did not identify any obvious 9aaTAD motif in the gained hyrax exon.

In the ray-finned bony fish *Danio rerio* (actinopterygians, see their low diversification in **Suppl. Figure S7**) and in the lobe-finned bony fish *Latimeria chalumnae* (sarcopterygians) we found a common modification of the *Sp2* activation domain (loss of valines in positions p1 and p8 by deletions at p1 and p2, and substitutions of p8). While the modifications are absent in the jawed cartilaginous fishes (chondrichthyes), the modifications must be ancestral for all bony fishes (osteichthyes) but not for all jawed fish (gnathostomes) (Figure 5). Both Sp2 became re-activated, which correlated with a loss of two out of three valines within their activation domains.

### The *Sp2 gene* duplication in *Takifugu* and *Tetraodon*

A fish-specific whole-genome duplication is present in some ray-finned fish lineages (actinopterygians) that branched from the sister lineage, lobe-finned fishes (sarcopterygians) about 450 Mya but is not present in terrestrial vertebrates (77). We have identified duplicated *Sp2 genes* (designated as *Sp2a* and *Sp2b*) in the teleost lineage of the ray-finned fishes takifugu and tetraodon, but not in other teleosts, such as *Danio rerio* (zebrafish) and *Oryzias latipes* (japanese medaka), nor in cartilaginous fish *Callorhinchus milii* (elephant shark), nor in lobe-finned fish *Latimeria chalumnae* (coelacanth), nor in terrestrial vertebrates (reptiles / alligator, birds / chicken, or mammals / opossum, mouse or human) (**Suppl. Figure S8**).

Both *Sp2* genes duplicated in takifugu and tetraodon lost their exons coding for their activation domains. These genes (*Sp2b genes*) were repaired from their intronal DNA, which generated new exons without obvious 9aaTAD activation domains. The origin of the new exons could be well recognized from their ancestral introns (present in takifugu and tetraodon *Sp2a genes*) (**Suppl. Figure S8**). The repairs occurred independently rather than originating from a common ancestor of the tetraodontiformes, because the *Sp2a* intronal sequences of takifugu and tetraodon lack sequence similarity. The relevant *Sp2* exon replacement could also be found in gibbon, in rock hyrax and in crane *Sp2* (Figure 7 and **Suppl. Figure S9**).

We have investigated the evolution of the new *Sp2* activation domains in detail and found that their intron sequences are reservoirs of activation domains. The introns have full readiness for activation domain repair by presence of i) a copy of exon-intron sequence border, which might promote DNA recombination in a desirable position, ii) a copy of the original 9aaTAD activation domain, which might be used as a backup for random loss of exonal activation domain and iii) repeats resembling the 9aaTAD domains in all three reading frames, which might facilitate *de novo* formation of another activation domain (the tested construct w35 including activation domain coded by **the tetraodon** intron, for which we have observed the capacity to activate transcription) (**Suppl. Figure S8**).

Although the *Sp2a gene* in tetraodon has two intronal activation domain back-ups ready to use (**Suppl. Figure S8**), these domains were not involved within the *Sp2b gene* rearrangement. We suspect that these events have been rather conditioned by evolutionary pressure to deactivate duplicated genes and not to use the activation domain from takifugu and tetraodon introns.

### The deactivated Sp2 turned into activator in gibbon

We have searched for losses of the original activation domains and their replacement by new sequences in *Sp2 genes.* We have identified new formations of the *Sp2 exon*, coding for activation domains in gibbon, in rock hyrax and in crane (Figure 5 and **Suppl. Figure S9**). Gibbons are “lesser apes” that belong to the anthropoid primates (hominoidea) superfamily together with the “great apes” (chimpanzees, gorillas, orangutans and humans). The *Sp2* exon, encoding a valine-rich activation domain is well conserved in all eutherian (placental mammals) including primates (Figure 5). Differently, the gibbon, rock hyrax and crane *Sp2* exons encoding the activation domain lack homology to the other eutherian *Sp2* exons and their activation domains are not valine-rich. Similar to takifugu, their *Sp2 genes* were broken at the same exon/intron junction (corresponding to the exon 3 and intron 3/4 junction in humans) and new exons have risen from their ancestral intron sequence. Therefore, the sequences encoding activation domains differ from those in other eutherian *Sp2 genes* (Figure 5).

The human *Sp2* intron 3/4 includes multiple putative activation domains in all three open reading frames (WQ-SWL, WQ-LFQV, SWQ-LLF, FWH-LFL, TWN-SW) with similarity to Sp1-4 clade activation domains (Sp1: SWQ-TLQL, Sp3: SWQ-TFQV, Sp4: SWQ-TVQV), that suggest the primates’ readiness for activation domain repair of the *Sp2 gene*, as we have already found in the tetraodon *Sp2a* intron. We used the human intron sequence for the reconstitution of the ancestral gibbon’s gene (Figure 5). We have identified the activation domain in the new gibbon exon 4, which has the 9aaTAD pattern and has no valines. We generated the construct w81 for the gibbon’s activation domain and tested the capacity to activate transcription. The rescued gibbon’s *Sp2 gene* encodes a new 9aaTAD activation domain, whose transcriptional activation capacity is comparable to the Gal4 activation domain (14). Gibbons have therefore recovered a strong activation domain encoded by the *Sp2 gene* that has not been seen since the cnidarians. The Sp2 activation domains were partially active in marsupials, but completely deactivated in placental mammals, including humans. Because of the new gibbon *Sp2 gene* with the strong activation domain becoming stable in gibbon population, we suspect that the repair was driven by evolutionary pressure for Sp2 activation function, which might be beneficial if not essential in gibbons.

In the gibbon’s case, the intron provides a new activation domain and the encoded protein regained activator function. From above, we found a similar *Sp2 gene* break and repair in another terrestrial mammal *Procavia capensis* (rock hyrax), which is a close relative of sirenians and elephants (clade atlantogenates). The *Sp2 gene* sequence was acquired from the UCSC genomic annotation for rock hyrax Sp2 scaffold_25:102095-130179. In this case, no obvious activation domain was found in their newly-generated exon. The hyrax *Sp2 gene* was broken and repaired on the border of exon 3 and exon 4, exactly as in takifugu, tetraodon, crane and gibbon. We conclude that conserved *Sp2* gene repairs, on border of the exon 3 and exon 4, are inborn.

### The use of intronal activation domains is not limited to the SP family

Similarly to the *Sp2 gene*, we have identified intronal 9aaTAD activation domains in the unrelated human SREBP1 gene, which alternatively engaged exon or intron coding for alternative activation domains. The SREBP1 exon 1 includes part of the activation domain 9aaTAD-I, which could be found only in isoforms 1, 2, 4 and 5. Another alternatively used SREBP1 exon2 includes part of the activation domain 9aaTAD-II, which could only be found in isoforms 3, 4 and 6. In summary, all isoforms of the human SREBP1 include one of the two possible activation domains (Figure 6).

**Figure 6.**
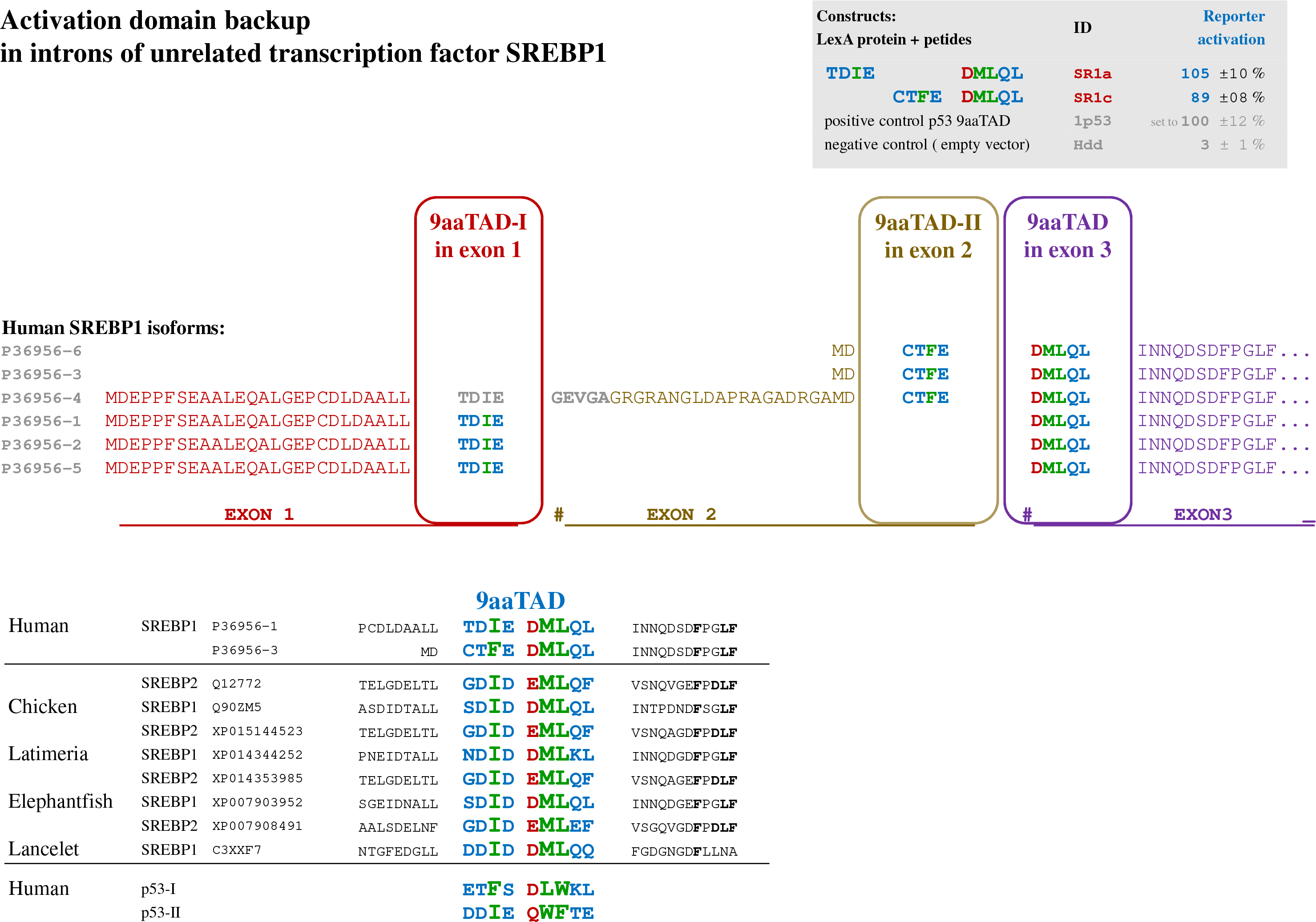
Activation domain backup in introns of unrelated transcription factor SREBP. The *exon 1* and *exon 2* junction to *exon 3* of *SREBP1 gene* coded for two distinct 9aaTAD activation domains (exon *1* in red, *exon 2* in sand brown, *exon 3* in purpura). The first activation domain 9aaTAD-I is coded by *exon 1* (used in isoform 1, 2 and 5 but not in 3, 4 and 6) and the second 9aaTAD-II coded by *exon 2* (used in isoform 3, 4 and 6 but not in 1, 2 and 5). The alternative splicing always resulted in the presence of a solitary 9aaTAD activation domain in all isoforms. The peptides with capacity to activate transcription are shown in the chart. The 9aaTAD motif conservation in the SREBP family and the similarity with p53 9aaTAD activation domains are shown in lower panel. The 9aaTAD activation domains are coloured for fast orientation. The exon borders are marked by coloured hashtags.

### KLF homology with the SP family revealed their activation domains

The SP clade is a new branch derived from the KLF family. We have reviewed similarities between the *SP* and *KLF genes* in various metazoans and have found the SP proteins diversified the least from the ancestral KLF proteins in <u>*S*</u>*trongylocentrotus* <u>pu</u>rpuratus (Spu, sea urchin) (Figure 7). The close similarity of the sea urchin *SP* and *KLF genes* enabled us to localise the 9aaTAD activation domains (from the SP and KLF sequence alignment in Figure 7). Next, by comparison of the sea urchin with the human *KLF genes* using simple BLAST search and BLAST generated alignments (not shown), we have identified the 9aaTADs in the human *KLF-1, 2, 3, 4, 5, 6, 7, 8, 12, 15* and *WT1 genes* (Wilms tumour protein).

**Figure 7.**
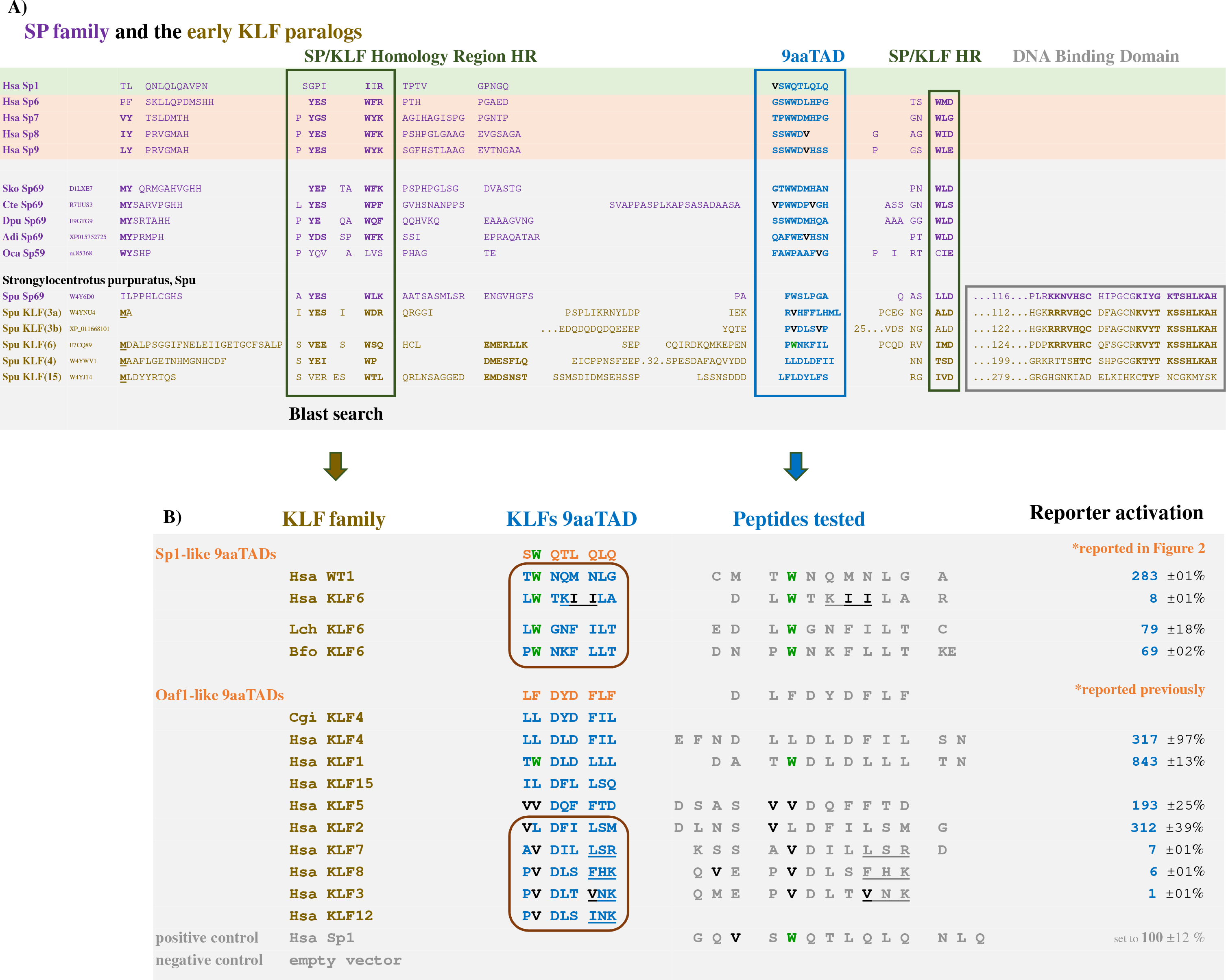
Alignment of activation sequences of *S.purpuratus* SP and KLF paralogs revealed activation domains in the KLF family. A) <u>*S*</u>*trongylocentrotus* <u>pu</u>rpuratus (sea urchin) <u>*Spu*</u>*Sp69* sequence was aligned to <u>*Spu*</u>KLF2/3/7/15 sequences (green box for homology region, blue box for 9aaTADs and black box for DNA binding domain). Human and other metazoan SP proteins are shown to have homology regions. The 9aaTAD activation domains in the human KLF proteins were identified by sequence similarity to the activation domain in <u>Spu</u>KLFs (simple BLAST search). B) Accordingly to sequence similarity, the KLF and WT1 9aaTAD activation domains were clustered into Sp1-like and Oaf1-like activation domain subgroups (DOI: 10.1016/j.ygeno.2007.02.003). * <u>*H*</u>omo sapiens Hsa, <u>*S*</u>*accoglossus* <u>ko</u>walevskii (acorn worm) <u>*Sko*</u>, <u>*C*</u>apitella <u>te</u>leta (annelid worm) <u>*Cte*</u>, <u>*D*</u>*aphnia* <u>pu</u>lex (water flea) <u>*Dpu*</u>, <u>*A*</u>*cropora* <u>*di*</u>*gitifera* (acroporid coral) <u>*Adi*</u>, <u>*O*</u>*scarella* <u>ca</u>rmela (slime sponge) <u>*Oca*</u>, <u>*C*</u>*rassostrea* <u>gi</u>gas (pacific oyster) <u>*Cgi*</u>), <u>*L*</u>*atimeria* <u>ch</u>alumnae (coelacanth) <u>*Lch*</u>, <u>*B*</u>*ranchiostoma* <u>flo</u>ridae (lancelet) <u>*Bfo*</u>. The valines and twin isoleucines (in <u>*Hsa*</u>*KLF6* only) are in black, phenylalanines in positions of valines (in <u>*Hsa*</u>*KLF3*) are in red and tryptophans are in green. Hidden amino acids are marked with points and numbers. Initial methionines are underlined. The regions with similarity of active and deactivated KLFs are in brown boxes.

Some of these identified activation domains correspond to previously reported activation domains. The activation domain of the KLF2 was first localised to its short N-terminal region (1–57) (78). Furthermore, the KLF4 activation domain has been characterised by loss-of-function mutations (EEE93/95/96VVV and DDD99/102/104VVV, in which both amino acid substitutions introduced valines in the KLF4 activation domain)(79). We have previously (12) identified the 9aaTAD motif in the KLF4 mutated region (9aaTAD position correlation in the SP and KLF families). Next, two activation domains were reported for the KLF1 (80), which are both also recognized by our 9aaTAD prediction algorithm (with one perfect match ATW <u>**DLD**</u> LLL) (12). Interestingly, we have found the 9aaTAD activation domain in *Crassostrea gigas, KLF1* ortholog (9aaTAD sequence ELL <u>**DYD**</u> FIL, gene ID; EKC39171), which closely resembles the first identified 9aaTAD in activator *Oaf1* (9aaTAD sequence DLF <u>**DYD**</u> FLF) (12).

Surprisingly, the 9aaTAD activation domains of KLF3 and KLF8 did not activate transcription, which correlated with their repressor function reported previously (52, 81–84). The accumulation of valines has been shown here again to be associated with the 9aaTAD deactivation (Figures 7). Nevertheless, the accumulation of valines in the KLF5 activation domain (in position p1 and p2 of the 9aaTAD) did not cause the 9aaTAD deactivation, therefore the accumulation of valines by itself is not always sufficient for 9aaTAD deactivation in any of its position.

Similarly to the human KLF6, in which the 9aaTAD domain has accumulated isoleucines, we did not find any capacity to activate transcription for this activation domain. Nevertheless, the evolutionary younger *KLF6 orthologs* in *Latimeria chalumnae* (Coelacanth, KLF6 gene ID; H3AK31) and *Branchiostoma floridae* (Florida lancelet, KLF6 gene ID; C3ZT77) have not accumulated isoleucines and hence their 9aaTAD activation domains have the capacity to activate transcription (Figures 8).

**Figure 8.**
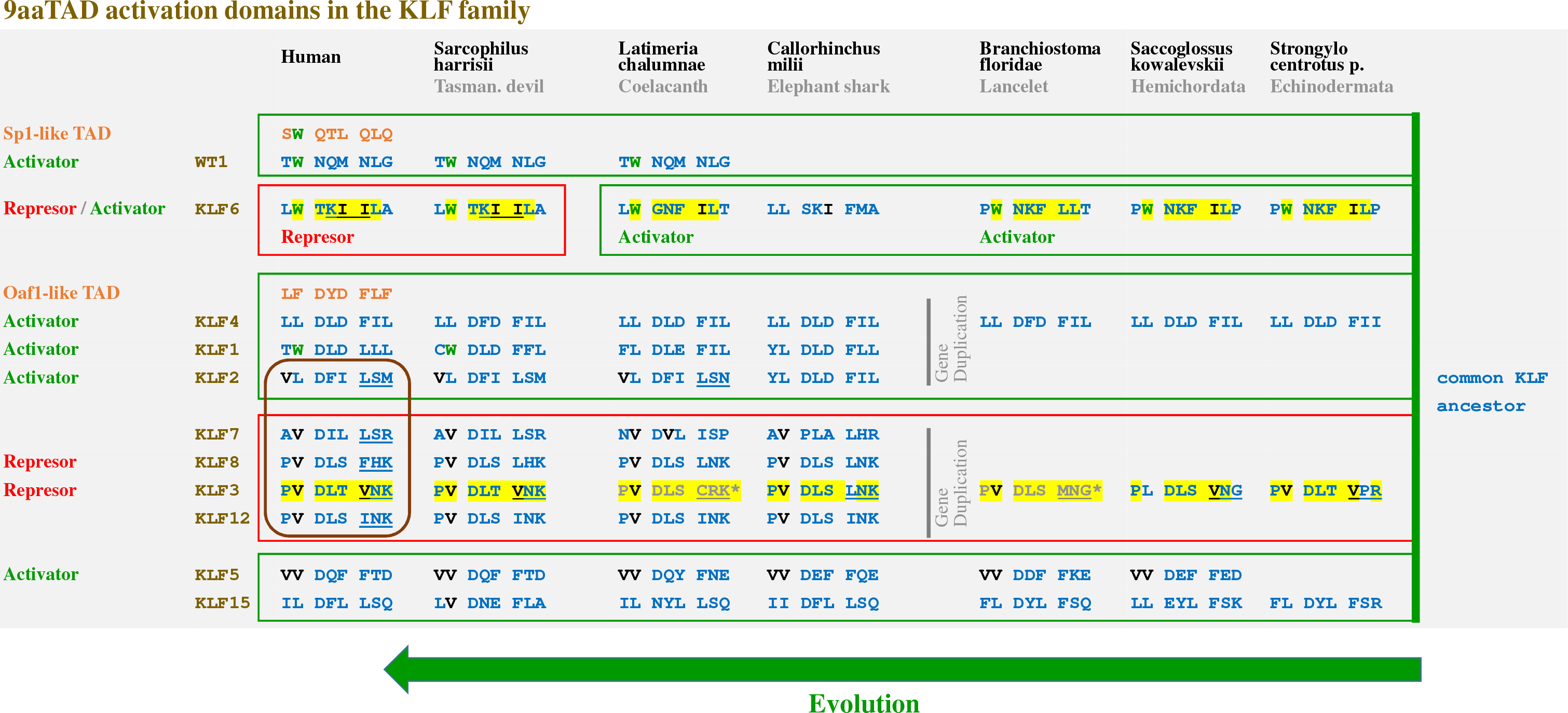
Schema of KLF evolution. The KLF genes from <u>*S*</u>*arcophilus* <u>har</u>risii (tasmanian devil) <u>*Shr*</u>, <u>L</u>atimeria <u>c</u>halu<u>m</u>nae (coelacanth) <u>*Lcm*</u>, <u>C</u>allorhinchus <u>m</u>ilii (elephant shark) <u>*Cmk*</u>, <u>B</u>ranchiostoma <u>flo</u>ridae (Florida lancelet) <u>*Bfo*</u>, <u>S</u>accoglossus <u>ko</u>walevskii (acorn worm) <u>*Sko*</u> and Amphimedon queenslandica are listed in Supp. Table T2. The designation numbers with brackets reflected uncertainty of origin. Sequences in blue fulfil the 9aaTAD pattern but sequences in grey and marked with an asterisk do not. The first 9aaTAD position p1 is the most variable, and therefore not shown in the similarity overview. The valines and isoleucines are in black and tryptophans are in green. * Repressor function for the KLF12 activation domain was only deduced from their reported repressor function and its sequence similarity to KLF3 and KLF6.

## DISCUSSION

Previously, we have defined a pattern for the 9aaTAD domains that we use in the algorithm in the online Prediction Tool (www.piskacek.org), which has allowed the identification of the activation domains in numerous transcriptional activators. In this study, we have identified the 9aaTAD domains in the glutamine-rich activator Sp1 and in the proline-rich activator NFIC/CTF. We have demonstrated that the aspartic acid and glutamine residues are interchangeable in the activation domains of the glutamine-rich activator Sp1 and in the acidic activator Gal4. Moreover, we have shown that the conservation of glutamines was limited to the SP1-4 clade in vertebrates, but was almost absent in protostomes and in poriferans. Although the SP6-9 clade shares sequence homology with the SP1-4 clade, they also did not accumulate glutamines in their activation domains. Consistently, the proline-rich activation domains of NFIC/CTF activators did not conserve prolines, but rather the 9aaTAD activation domain, through the family. Thus, we united these previously classified acidic, glutamine-rich and proline-rich activators into the 9aaTAD family.

Next, we have described correlation of the 9aaTAD deactivation with accumulation of valines (in human Sp2 and KLF3), leucines (in *Latimeria* and in marsupial Sp2) and isoleucines (in human KLF6). The branched valine or isoleucine amino acids could partially hinder formation of α-helixes and cause their destabilization (85). Therefore, the accumulation of valines and isoleucines, which do not fully support helix formation, might partially explain the 9aaTAD deactivation. Nevertheless, the accumulation of valines at the end of the KLF5 activation domain (in position p1 and p2 of the 9aaTAD) did not cause the 9aaTAD deactivation, therefore the accumulation of valines itself in any position of the 9aaTAD (especially in position p1 and p2 in the KLF5 or in position p3 in Gcn4 or Gal4), is not always sufficient for the deactivation. Similarly, we have identified the accumulation of valines in the 9aaTAD activation domain in a strong activator Met4, which again did not interfere with the activation function (Met4 activation domain was found by the sequence similarity with the deactivated *Sp2 genes*; ScanProsite search (ExPASy) for V-[NQDE]-W-[QN] amino acid pattern) (**Suppl. Figure S10**). From the limited number of activation domains with accumulated valines, we could not yet elucidate the amino acid pattern, which is necessary for deactivation of the 9aaTAD domain.

Importantly, we have observed re-activation of the 9aaTAD activation domains after their *de-valinisation* in the SP family. Similarly to the SP family, *de-valinisation* of the Gcn4 activation domain by substitution with tryptophans has been reported as a great enhancement of the otherwise weak Gcn4 activation domain (A<u>**VV**</u>E SFFSS → A<u>**V**</u>wE SLFSS → A**www w**LF**w**S) (86). Additionally, we have previously shown a partial *de-valinisation* of the Gcn4 activation domain by valine substitution for aspartic acid with similar results to increase its function (A<u>**VV**</u>E SFFSS → DD<u>**V**</u>E SLFFS → DD<u>**V**</u>Y NYLFD) (14).

In the gibbon, we found surprising recovery of the damaged *Sp2 gene* that has reverted into an activator, the very opposite function of the original gene, otherwise found in all other primates. Such dramatic changes in gene regulation in primates were highly unexpected. Nevertheless, the *Sp2 genes* are stable in gibbon population and might be a part of their accelerated genome evolution (87). Mouse knock-ins of the gibbon *Sp2 gene* coding for an activator might clarify if the Sp2 re-activation could be beneficial for other mammals or would have driven gibbon accelerated evolution.

The most intriguing observation in this study has been the preservation of the deactivated Sp2 9aaTAD domains, which lasted over 100 million of years. After the *Sp2 gene* duplication in primitive fish (jawless and cartilaginous fish), the new paralogs were able to diversify into both activators and repressors and have provided a larger diversification of the SP regulatory system (88). After the *Sp2 gene* duplication, the newly born paralog *Sp1* has regained its activator function by *de-valinisation* of the 9aaTAD activation domain. The domain deactivation, rather than simple domain deletion, was obviously an advantage during vertebrate evolution.

## References

1. Teufel,D.P., Freund,S.M., Bycroft,M. and Fersht,A.R. (2007) Four domains of p300 each bind tightly to a sequence spanning both transactivation subdomains of p53. Proc. Natl. Acad. Sci. U.S.A., 104, 7009–7014.

2. Gamper,A.M. and Roeder,R.G. (2008) Multivalent Binding of p53 to the STAGA Complex Mediates Coactivator Recruitment after UV Damage. Mol Cell Biol, 28, 2517–2527.

3. Feng,H., Jenkins,LM.M., Durell,S.R., Hayashi,R., Mazur,S.J., Cherry,S., Tropea,J.E., Miller,M., Wlodawer,A., Appella,E., et al. (2009) Structural basis for p300 Taz2-p53 TAD1 binding and modulation by phosphorylation. Structure, 17, 202–210.

4. Ferreon,J.C., Lee,C.W., Arai,M., Martinez-Yamout,M.A., Dyson,H.J. and Wright,P.E. (2009) Cooperative regulation of p53 by modulation of ternary complex formation with CBP/p300 and HDM2. Proc. Natl. Acad. Sci. U.S.A., 106, 6591–6596.

5. Jenkins,LM.M., Yamaguchi,H., Hayashi,R., Cherry,S., Tropea,J.E., Miller,M., Wlodawer,A., Appella,E. and Mazur,S.J. (2009) Two distinct motifs within the p53 transactivation domain bind to the Taz2 domain of p300 and are differentially affected by phosphorylation. Biochemistry, 48, 1244–1255.

6. Thakur,J.K., Arthanari,H., Yang,F., Chau,K.H., Wagner,G. and Näär,A.M. (2009) Mediator subunit Gal11p/MED15 is required for fatty acid-dependent gene activation by yeast transcription factor Oaf1p. J. Biol. Chem., 284, 4422–4428.

7. Choi,Y., Asada,S. and Uesugi,M. (2000) Divergent hTAFII31-binding motifs hidden in activation domains. J. Biol. Chem., 275, 15912–15916.

8. Uesugi,M. and Verdine,G.L. (1999) The alpha-helical FXXPhiPhi motif in p53: TAF interaction and discrimination by MDM2. Proc. Natl. Acad. Sci. U.S.A., 96, 14801–14806.

9. Piskacek,M. (2009) 9aaTADs mimic DNA to interact with a pseudo-DNA Binding Domain KIX of Med15 (Molecular Chameleons). Nature Precedings, 10.1038/npre.2009.3939.1.

10. Piskacek,M. (2009) Common Transactivation Motif 9aaTAD recruits multiple general co-activators TAF9, MED15, CBP and p300. Nature Precedings, 10.1038/npre.2009.3488.2.

11. Di Lello,P., Jenkins,LM.M., Jones,T.N., Nguyen,B.D., Hara,T., Yamaguchi,H., Dikeakos,J.D., Appella,E., Legault,P. and Omichinski,J.G. (2006) Structure of the Tfb1/p53 complex: Insights into the interaction between the p62/Tfb1 subunit of TFIIH and the activation domain of p53. Mol. Cell, 22, 731–740.

12. Piskacek,S., Gregor,M., Nemethova,M., Grabner,M., Kovarik,P. and Piskacek,M. (2007) Nine-amino-acid transactivation domain: establishment and prediction utilities. Genomics, 89, 756–768.

13. Piskacek,M., Vasku,A., Hajek,R. and Knight,A. (2015) Shared structural features of the 9aaTAD family in complex with CBP. Mol Biosyst, 11, 844–851.

14. Piskacek,M., Havelka,M., Rezacova,M. and Knight,A. (2016) The 9aaTAD Transactivation Domains: From Gal4 to p53. PLoS ONE, 11, e0162842.

15. Piskacek,M. (2009) 9aaTAD Prediction result (2006). Nature Precedings, 10.1038/npre.2009.3984.1.

16. Sandholzer,J., Hoeth,M., Piskacek,M., Mayer,H. and de Martin,R. (2007) A novel 9-amino-acid transactivation domain in the C-terminal part of Sox18. Biochem. Biophys. Res. Commun., 360, 370–374.

17. Piskacek,M., Havelka,M., Rezacova,M. and Knight,A. (2017) The 9aaTAD Is Exclusive Activation Domain in Gal4. PLoS ONE, 12, e0169261.

18. Kakidani,H. and Ptashne,M. (1988) GAL4 activates gene expression in mammalian cells. Cell, 52, 161–167.

19. Fields,S. and Jang,S.K. (1990) Presence of a potent transcription activating sequence in the p53 protein. Science, 249, 1046–1049.

20. Triezenberg,S.J. (1995) Structure and function of transcriptional activation domains. Curr. Opin. Genet. Dev., 5, 190–196.

21. Ma,J. and Ptashne,M. (1987) A new class of yeast transcriptional activators. Cell, 51, 113–119.

22. Courey,A.J. and Tjian,R. (1988) Analysis of Sp1 in vivo reveals multiple transcriptional domains, including a novel glutamine-rich activation motif. Cell, 55, 887–898.

23. Courey,A.J., Holtzman,D.A., Jackson,S.P. and Tjian,R. (1989) Synergistic activation by the glutamine-rich domains of human transcription factor Sp1. Cell, 59, 827–836.

24. Tanese,N., Pugh,B.F. and Tjian,R. (1991) Coactivators for a proline-rich activator purified from the multisubunit human TFIID complex. Genes Dev., 5, 2212–2224.

25. Mermod,N., O’Neill,E.A., Kelly,T.J. and Tjian,R. (1989) The proline-rich transcriptional activator of CTF/NF-I is distinct from the replication and DNA binding domain. Cell, 58, 741–753.

26. Stargell,L.A. and Struhl,K. (1995) The TBP-TFIIA interaction in the response to acidic activators in vivo. Science, 269, 75–78.

27. Chou,S. and Struhl,K. (1997) Transcriptional activation by TFIIB mutants that are severely impaired in interaction with promoter DNA and acidic activation domains. Mol. Cell. Biol., 17, 6794–6802.

28. Dorris,D.R. and Struhl,K. (2000) Artificial Recruitment of TFIID, but Not RNA Polymerase II Holoenzyme, Activates Transcription in Mammalian Cells. Mol Cell Biol, 20, 4350–4358.

29. Thoden,J.B., Ryan,L.A., Reece,R.J. and Holden,H.M. (2008) The interaction between an acidic transcriptional activator and its inhibitor. The molecular basis of Gal4p recognition by Gal80p. J. Biol. Chem., 283, 30266–30272.

30. Drysdale,C.M., Dueñas,E., Jackson,B.M., Reusser,U., Braus,G.H. and Hinnebusch,A.G. (1995) The transcriptional activator GCN4 contains multiple activation domains that are critically dependent on hydrophobic amino acids. Mol. Cell. Biol., 15, 1220–1233.

31. Jackson,B.M., Drysdale,C.M., Natarajan,K. and Hinnebusch,A.G. (1996) Identification of seven hydrophobic clusters in GCN4 making redundant contributions to transcriptional activation. Mol. Cell. Biol., 16, 5557–5571.

32. Natarajan,K., Meyer,M.R., Jackson,B.M., Slade,D., Roberts,C., Hinnebusch,A.G. and Marton,M.J. (2001) Transcriptional profiling shows that Gcn4p is a master regulator of gene expression during amino acid starvation in yeast. Mol. Cell. Biol., 21, 4347–4368.

33. Jedidi,I., Zhang,F., Qiu,H., Stahl,S.J., Palmer,I., Kaufman,J.D., Nadaud,P.S., Mukherjee,S., Wingfield,P.T., Jaroniec,C.P., et al. (2010) Activator Gcn4 employs multiple segments of Med15/Gal11, including the KIX domain, to recruit mediator to target genes in vivo. J. Biol. Chem., 285, 2438–2455.

34. Krois,A.S., Ferreon,J.C., Martinez-Yamout,M.A., Dyson,H.J. and Wright,P.E. (2016) Recognition of the disordered p53 transactivation domain by the transcriptional adapter zinc finger domains of CREB-binding protein. Proc. Natl. Acad. Sci. U.S.A., 113, E1853–1862.

35. Lee,C.W., Arai,M., Martinez-Yamout,M.A., Dyson,H.J. and Wright,P.E. (2009) Mapping the interactions of the p53 transactivation domain with the KIX domain of CBP. Biochemistry, 48, 2115–2124.

36. Denis,C.M., Chitayat,S., Plevin,M.J., Wang,F., Thompson,P., Li,S., Spencer,H.L., Ikura,M., Lebrun,D.P. and Smith,S.P. (2012) Structural basis of CBP/p300 recruitment in leukemia induction by E2A-PBX1. Blood, 10.1182/blood-2012-02-411397.

37. Wang,F., Marshall,C.B., Li,G.-Y., Yamamoto,K., Mak,T.W. and Ikura,M. (2009) Synergistic interplay between promoter recognition and CBP/p300 coactivator recruitment by FOXO3a. ACS Chem. Biol., 4, 1017–1027.

38. Radhakrishnan,I., Pérez-Alvarado,G.C., Parker,D., Dyson,H.J., Montminy,M.R. and Wright,P.E. (1997) Solution structure of the KIX domain of CBP bound to the transactivation domain of CREB: a model for activator:coactivator interactions. Cell, 91, 741–752.

39. Lee,C.W., Martinez-Yamout,M.A., Dyson,H.J. and Wright,P.E. (2010) Structure of the p53 transactivation domain in complex with the nuclear receptor coactivator binding domain of CREB binding protein. Biochemistry, 49, 9964–9971.

40. Wojciak,J.M., Martinez-Yamout,M.A., Dyson,H.J. and Wright,P.E. (2009) Structural basis for recruitment of CBP/p300 coactivators by STAT1 and STAT2 transactivation domains. EMBO J., 28, 948–958.

41. Gill,G., Pascal,E., Tseng,Z.H. and Tjian,R. (1994) A glutamine-rich hydrophobic patch in transcription factor Sp1 contacts the dTAFII110 component of the Drosophila TFIID complex and mediates transcriptional activation. Proc. Natl. Acad. Sci. U.S.A., 91, 192–196.

42. Titz,B., Thomas,S., Rajagopala,S.V., Chiba,T., Ito,T. and Uetz,P. (2006) Transcriptional activators in yeast. Nucleic Acids Res., 34, 955–967.

43. Escher,D., Bodmer-Glavas,M., Barberis,A. and Schaffner,W. (2000) Conservation of glutamine-rich transactivation function between yeast and humans. Mol. Cell. Biol., 20, 2774–2782.

44. Hahn,S. (1993) Structure(?) and function of acidic transcription activators. Cell, 72, 481–483.

45. Brzovic,P.S., Heikaus,C.C., Kisselev,L., Vernon,R., Herbig,E., Pacheco,D., Warfield,L., Littlefield,P., Baker,D., Klevit,R.E., et al. (2011) The acidic transcription activator Gcn4 binds the mediator subunit Gal11/Med15 using a simple protein interface forming a fuzzy complex. Mol. Cell, 44, 942–953.

46. Lu,Z., Ansari,A.Z., Lu,X., Ogirala,A. and Ptashne,M. (2002) A target essential for the activity of a nonacidic yeast transcriptional activator. Proc. Natl. Acad. Sci. U.S.A., 99, 8591–8596.

47. Ma,J. and Ptashne,M. (1987) Deletion analysis of GAL4 defines two transcriptional activating segments. Cell, 48, 847–853.

48. Ferreira,M.E., Hermann,S., Prochasson,P., Workman,J.L., Berndt,K.D. and Wright,A.P.H. (2005) Mechanism of transcription factor recruitment by acidic activators. J. Biol. Chem., 280, 21779–21784.

49. Staller,M.E., Holehouse,A.S., Swain-Lenz,D., Das,R.K., Pappu,R.V. and Cohen,B.A. (2018) A High-Throughput Mutational Scan of an Intrinsically Disordered Acidic Transcriptional Activation Domain. Cell Syst, 10.1016/j.cels.2018.01.015.

50. Zhang,H.-M., Liu,T., Liu,C.-J., Song,S., Zhang,X., Liu,W., Jia,H., Xue,Y. and Guo,A.-Y. (2015) AnimalTFDB 2.0: a resource for expression, prediction and functional study of animal transcription factors. Nucleic Acids Res., 43, D76–81.

51. Kolell,K.J. and Crawford,D.L. (2002) Evolution of Sp transcription factors. Mol. Biol. Evol., 19, 216–222.

52. Kaczynski,J., Cook,T. and Urrutia,R. (2003) Sp1-and Krüppel-like transcription factors. Genome Biol., 4, 206.

53. Suske,G., Bruford,E. and Philipsen,S. (2005) Mammalian SP/KLF transcription factors: bring in the family. Genomics, 85, 551–556.

54. Vizcaíno,C., Mansilla,S. and Portugal,J. (2015) Sp1 transcription factor: A long-standing target in cancer chemotherapy. Pharmacol. Ther., 152, 111–124.

55. Mir,R., Sharma,A., Pradhan,S.J. and Galande,S. (2018) Regulation of transcription factor SP1 by β-catenin destruction complex modulates Wnt response. bioRxiv, 10.1101/308841.

56. Rane,M.J., Zhao,Y. and Cai,L. (2019) Krϋppel-like factors (KLFs) in renal physiology and disease. EBioMedicine, 10.1016/j.ebiom.2019.01.021.

57. Miller,J.H. (1972) Experiments in molecular genetics Cold Spring Harbor Laboratory.

58. Baumgartner,U., Hamilton,B., Piskacek,M., Ruis,H. and Rottensteiner,H. (1999) Functional analysis of the Zn(2)Cys(6) transcription factors Oaf1p and Pip2p. Different roles in fatty acid induction of beta-oxidation in Saccharomyces cerevisiae. J. Biol. Chem., 274, 22208–22216.

59. Leuther,K.K., Salmeron,J.M. and Johnston,S.A. (1993) Genetic evidence that an activation domain of GAL4 does not require acidity and may form a beta sheet. Cell, 72, 575–585.

60. Baur,F., Nau,K., Sadic,D., Allweiss,L., Elsässer,H.-P., Gillemans,N., de Wit,T., Krüger,I., Vollmer,M., Philipsen,S., et al. (2010) Specificity protein 2 (Sp2) is essential for mouse development and autonomous proliferation of mouse embryonic fibroblasts. PLoS ONE, 5, e9587.

61. Terrados,G., Finkernagel,F., Stielow,B., Sadic,D., Neubert,J., Herdt,O., Krause,M., Scharfe,M., Jarek,M. and Suske,G. (2012) Genome-wide localization and expression profiling establish Sp2 as a sequence-specific transcription factor regulating vitally important genes. Nucleic Acids Res., 40, 7844–7857.

62. Völkel,S., Stielow,B., Finkernagel,F., Stiewe,T., Nist,A. and Suske,G. (2015) Zinc finger independent genome-wide binding of Sp2 potentiates recruitment of histone-fold protein Nf-y distinguishing it from Sp1 and Sp3. PLoS Genet., 11, e1005102.

63. Ratajewski,M., Walczak-Drzewiecka,A., Gorzkiewicz,M., Sałkowska,A. and Dastych,J. (2016) Expression of human gene coding RORγT receptor depends on the Sp2 transcription factor. J. Leukoc. Biol., 100, 1213–1223.

64. Zschemisch,N.-H., Brüsch,I., Hambusch,A.-S. and Bleich,A. (2016) Transcription Factor SP2 Enhanced the Expression of Cd14 in Colitis-Susceptible C3H/HeJBir. PLoS ONE, 11, e0155821.

65. Moorefield,K.S., Fry,S.J. and Horowitz,J.M. (2004) Sp2 DNA binding activity and trans-activation are negatively regulated in mammalian cells. J. Biol. Chem., 279, 13911–13924.

66. Yin,H., Nichols,T.D. and Horowitz,J.M. (2010) Transcription of mouse Sp2 yields alternatively spliced and sub-genomic mRNAs in a tissue-and cell-type-specific fashion. Biochim. Biophys. Acta, 1799, 520–531.

67. Phan,D., Cheng,C.-J., Galfione,M., Vakar-Lopez,F., Tunstead,J., Thompson,N.E., Burgess,R.R., Najjar,S.M., Yu-Lee,L.-Y. and Lin,S.-H. (2004) Identification of Sp2 as a transcriptional repressor of carcinoembryonic antigen-related cell adhesion molecule 1 in tumorigenesis. Cancer Res., 64, 3072–3078.

68. Yesudhas,D., Anwar,M.A., Panneerselvam,S., Kim,H.-K. and Choi,S. (2017) Evaluation of Sox2 binding affinities for distinct DNA patterns using steered molecular dynamics simulation. FEBS Open Bio, 7, 1750–1767.

69. Kamachi,Y. and Kondoh,H. (2013) Sox proteins: regulators of cell fate specification and differentiation. Development, 140, 4129–4144.

70. Lodato,M.A., Ng,C.W., Wamstad,J.A., Cheng,A.W., Thai,K.K., Fraenkel,E., Jaenisch,R. and Boyer,L.A. (2013) SOX2 co-occupies distal enhancer elements with distinct POU factors in ESCs and NPCs to specify cell state. PLoS Genet., 9, e1003288.

71. Ward,S.V. and Samuel,C.E. (2003) The pkr kinase promoter binds both Sp1 and Sp3, but only Sp3 functions as part of the interferon-inducible complex with ISGF-3 proteins. Virology, 313, 553–566.

72. Jaiswal,A.S., Balusu,R. and Narayan,S. (2006) 7,12-Dimethylbenzanthracene-dependent transcriptional regulation of adenomatous polyposis coli (APC) gene expression in normal breast epithelial cells is mediated by GC-box binding protein Sp3. Carcinogenesis, 27, 252–261.

73. Li,L. and Davie,J.R. (2010) The role of Sp1 and Sp3 in normal and cancer cell biology. Annals of Anatomy-Anatomischer Anzeiger, 192, 275–283.

74. Erwin,D.H., Laflamme,M., Tweedt,S.M., Sperling,E.A., Pisani,D. and Peterson,K.J. (2011) The Cambrian conundrum: early divergence and later ecological success in the early history of animals. Science, 334, 1091–1097.

75. Presnell,J.S., Schnitzler,C.E. and Browne,W.E. (2015) KLF/SP Transcription Factor Family Evolution: Expansion, Diversification, and Innovation in Eukaryotes. Genome Biol Evol, 7, 2289–2309.

76. Hackett,S.J., Kimball,R.T., Reddy,S., Bowie,R.C.K., Braun,E.L., Braun,M.J., Chojnowski,J.L., Cox,W.A., Han,K.-L., Harshman,J., et al. (2008) A phylogenomic study of birds reveals their evolutionary history. Science, 320, 1763–1768.

77. Christoffels,A., Koh,E.G.L., Chia,J.-M., Brenner,S., Aparicio,S. and Venkatesh,B. (2004) Fugu genome analysis provides evidence for a whole-genome duplication early during the evolution of ray-finned fishes. Mol. Biol. Evol., 21, 1146–1151.

78. Conkright,M.D., Wani,M.A. and Lingrel,J.B. (2001) Lung Krüppel-like factor contains an autoinhibitory domain that regulates its transcriptional activation by binding WWP1, an E3 ubiquitin ligase. J. Biol. Chem., 276, 29299–29306.

79. Geiman,D.E., Ton-That,H., Johnson,J.M. and Yang,V.W. (2000) Transactivation and growth suppression by the gut-enriched Krüppel-like factor (Krüppel-like factor 4) are dependent on acidic amino acid residues and protein-protein interaction. Nucleic Acids Res., 28, 1106–1113.

80. Mas,C., Lussier-Price,M., Soni,S., Morse,T., Arseneault,G., Di Lello,P., Lafrance-Vanasse,J., Bieker,J.J. and Omichinski,J.G. (2011) Structural and functional characterization of an atypical activation domain in erythroid Kruppel-like factor (EKLF). Proc. Natl. Acad. Sci. U.S.A., 108, 10484–10489.

81. Knights,A.J., Yik,J.J., Mat Jusoh,H., Norton,L.J., Funnell,A.P.W., Pearson,R.C.M., Bell-Anderson,K.S., Crossley,M. and Quinlan,K.G.R. (2016) Krüppel-like Factor 3 (KLF3/BKLF) Is Required for Widespread Repression of the Inflammatory Modulator Galectin-3 (Lgals3). J. Biol. Chem., 291, 16048–16058.

82. Klein,R.H., Hu,W., Kashgari,G., Lin,Z., Nguyen,T., Doan,M. and Andersen,B. (2017) Characterization of enhancers and the role of the transcription factor KLF7 in regulating corneal epithelial differentiation. J. Biol. Chem., 292, 18937–18950.

83. Das,A., Fernandez-Zapico,M.E., Cao,S., Yao,J., Fiorucci,S., Hebbel,R.P., Urrutia,R. and Shah,V.H. (2006) Disruption of an SP2/KLF6 repression complex by SHP is required for farnesoid X receptor-induced endothelial cell migration. J. Biol. Chem., 281, 39105–39113.

84. Zhang,H., Zhu,X., Chen,J., Jiang,Y., Zhang,Q., Kong,C., Xing,J., Ding,L., Diao,Z., Zhen,X., et al. (2015) Krüppel-like factor 12 is a novel negative regulator of forkhead box O1 expression: a potential role in impaired decidualization. Reprod. Biol. Endocrinol., 13, 80.

85. Pace,C.N. and Scholtz,J.M. (1998) A helix propensity scale based on experimental studies of peptides and proteins. Biophys. J., 75, 422–427.

86. Warfield,L., Tuttle,L.M., Pacheco,D., Klevit,R.E. and Hahn,S. (2014) A sequence-specific transcription activator motif and powerful synthetic variants that bind Mediator using a fuzzy protein interface. Proc. Natl. Acad. Sci. U.S.A., 111, E3506–3513.

87. Carbone,L., Harris,R.A., Gnerre,S., Veeramah,K.R., Lorente-Galdos,B., Huddleston,J., Meyer,T.J., Herrero,J., Roos,C., Aken,B., et al. (2014) Gibbon genome and the fast karyotype evolution of small apes. Nature, 513, 195–201.

88. Chong,S., Dugast-Darzacq,C., Liu,Z., Dong,P., Dailey,G.M., Cattoglio,C., Heckert,A., Banala,S., Lavis,L., Darzacq,X., et al. (2018) Imaging dynamic and selective low-complexity domain interactions that control gene transcription. Science, 361.

